# *In vitro* generation of megakaryocytes from engineered mouse embryonic stem cells

**DOI:** 10.1101/2023.03.01.530476

**Authors:** Mitchell R. Lewis, Tara L. Deans

**Affiliations:** Department of Biomedical Engineering, University of Utah, Salt Lake City, UT 84112, USA

## Abstract

The *in vitro* differentiation of pluripotent stem cells into desired lineages enables mechanistic studies of cell transitions into more mature states that can provide insights into the design principles governing cell fate control. We are interested in reprogramming pluripotent stem cells with synthetic gene circuits to drive mouse embryonic stem cells (mESCs) down the hematopoietic lineage for the production of megakaryocytes, the progenitor cells for platelets. Here, we describe the methodology for growing and differentiating mESCs, in addition to inserting a transgene to observe its expression throughout differentiation. This entails four key methods: (1) growing and preparing mouse embryonic fibroblasts for supporting mESC growth and expansion, (2) growing and preparing OP9 feeder cells to support the differentiation of mESCs, (3) the differentiation of mESCs into megakaryocytes, and (4) utilizing an integrase mediated docking site to insert transgenes for their stable integration and expression throughout differentiation. Altogether, this approach demonstrates a streamline differentiation protocol that emphasizes the reprogramming potential of mESCs that can be used for future mechanistic and therapeutic studies of controlling cell fate outcomes.

## 1. Introduction

Pluripotent stem cells have the potential to produce the three primary germ layers that make up the mammalian body. During development, embryonic stem cells (ESCs) undergo specialized decisionmaking to yield tissue specific characteristics that allow them to perform particular functions. Understanding how these cells tightly control their spatial and temporal gene expression of lineagespecific transcription factors will provide new insights into the design principles governing how cells transition from one cell state to another. Megakaryocytes (MKs) are a rare population of cells that develop from hematopoietic stem cells (HSCs) in the bone marrow and function to produce platelets that circulate throughout the body [1]. The misguided differentiation cues and incomplete maturation of MKs have been shown to cause a number of blood disorders [2]. In a healthy individual, roughly 1 in 10,000 bone marrow cells are MKs, making their development and maturation difficult to study. Another major challenge in studying MK development has been the identification, classification, and enrichment of MK progenitor cells that are produced during hematopoiesis, the process of making all cells of the blood system. We and others have recently identified, isolated, and expanded an important MK progenitor cell population that marks a critical transition state for stem cells in their journey to become mature MKs for the production of platelets [3–5]. In order to study the cell fate transitions and establish the molecular rules governing hematopoietic commitment, cell fate transitions, and the dynamic processes guiding stem cell differentiation during the development and maturation of MKs, it is critical to have a consistent and well-established protocol for the differentiation of pluripotent stem cells into MKs. Mouse embryonic stem cells (mESCs) are particularly interesting because of the large number of mouse disease models that exist for future studies of abnormal MK development and maturation, in addition to testing the function of platelets made *in vitro* [6] for mitigating disease, and during infection.

Here, we describe a new approach for differentiating mESCs *in vitro* for the production of MKs. Indeed, protocols exist in the literature for differentiating mESCs into MKs *in vitro*, however, many protocols contradict each other, give inconsistent results, and their MK production is minimal. Taking what we learned from attempting many protocols, we have established a new protocol for the consistent and robust differentiation of mESCs into MKs *in vitro*. Next, we show that mESCs can be engineered to express exogenous transgenes throughout differentiation, demonstrating the potential for reprogramming pluripotent stem cells with exogenous transgenes and synthetic gene circuits [7–13] to be used for driving their cell fate. In this protocol we describe differentiating mESCs on a support layer of OP9 cells, a bone marrow derived stromal cell line, in the presence of thrombopoietin (TPO) that leads to the *in vitro* differentiation of mESCs into MKs. We also describe how to use phage integrases to enable site-specific genome editing to insert desired DNA into the genome. Specifically, we use φC31 integrase, which catalyzes the irreversible recombination between appropriate *attB* and *attP* sites to insert desired DNA sequences [14, 15]. To accomplish this, we use mESCs from the TARGATT mouse line that have an *attP* site integrated in the Hipp11 chromosome [16]. We transfected mESCs with plasmids containing the φC31 integrase and GFP with *attB* sites flanking a promoter and GFP to enable its integration into the Hipp11 chromosome. After confirming integration, these mESCs were differentiated into MKs. Using the protocols outlined here will enable the growth and expansion of mESCs, the differentiation of mESCs into MKs *in vitro*, and studies for inserting transgenes and synthetic gene circuits into the Hipp11 chromosome of mESCs. These methods will facilitate studying the molecular rules and dynamic processes governing cell fate transitions into MKs, in addition to the mechanisms of transgene silencing, the challenge of losing expression of the inserted transgene(s) over time [17, 18].

## 2. Materials

### 2.1 Expanding and mitomycin C treating mouse embryonic fibroblasts (MEFs)

1. Primary mouse embryonic fibroblasts, neomycin resistant, not treated, P3 (Millipore Sigma, PMEF-NL).
2. MEF growth medium: 500 mL high glucose Dulbecco’s Modified Eagle’s Medium (DMEM, Thermo Fisher Scientific, 11965-092), 50 mL Fetal bovine serum (FBS; 10% final concentration, Thermo Fisher Scientific, 10437028), 5 mL penicillin/streptomycin (1% final concentration, Thermo Fisher Scientific, 15140-122), 5 mL L-glutamine (2mM final concentration, Thermo Fisher Scientific, 25030-081), 5 mL non-essential amino acids (NEAA; 1X final concentration, Thermo Fisher Scientific, 11140050).
3. Gelatin (Sigma, G1890).
4. Mitomycin C (MMC) (Fisher Scientific, BP2531-2).
5. DMSO (Fisher Scientific, MT-25950CQC).
6. 0.25% Trypsin-EDTA, phenol red (Thermo Fisher Scientific, 25200056).
7. PBS (Thermo Fisher Scientific, 10010-023).
8. MilliQ water.
9. T75 flasks, vented cap (Thermo Fisher Scientific, 12-556-010).
10. T175 flasks, vented cap (Thermo Fisher Scientific, 12-556-011).
11. 15 mL conical tubes (Fisher Scientific, 553912).
12. 50 mL conical tubes (Fisher Scientific, 05-539-8).
13. Sterile disposable filter unit, 250 mL (Fisher Scientific, FB12566502).
14. 70% ethanol in a spray bottle.
15. Centrifuge.
16. Hemocytometer.
17. Cryovials (Fisher Scientific, 03-337-7C).
18. Magnetic stir bar.
19. Magnetic stirrer hot plate.
20. Cell culture incubator capable of regulating temperature (37°C), humidity, and carbon dioxide (5%).
21. Biosafety hood suitable for growing cells aseptically.

### 2.2 Expanding mouse embryonic stem cells (mESCs)

1. Mouse embryonic stem cells (harvested from a TARGATT mouse, Applied StemCell) (*see* **Note 1**).
2. Confluent T25 flask of mitomycin C treated MEFs
3. mESC growth medium: 500 mL high glucose knock-out Dulbecco’s Modified Eagle’s Medium (DMEM, Thermo Fisher Scientific 10829-018), 75 mL fetal bovine serum (FBS), embryonic stem cell certified (FBS, ES certified; 15% final concentration, Thermo Fisher Scientific, 10439024), 5 mL penicillin/streptomycin (1% final concentration, Thermo Fisher Scientific, 15140-122), 5 mL L-glutamine (2mM final concentration, Thermo Fisher Scientific, 25030-081), 5 mL non-essential amino acids (NEAA; 1X final concentration, Thermo Fisher Scientific, 11140050), 500 μL β-mercaptoethanol (21985-023), Leukemia Inhibitory Factor (LIF, Thermo Fisher Scientific, PMC9484, *see* **Note 2**).
4. 0.25% Trypsin-EDTA, phenol red (Thermo Fisher Scientific, 25200056).
5. T25 flasks, vented cap (Thermo Fisher Scientific, 12556009).
6. 15 mL conical tubes (Fisher Scientific, 553912).
7. Centrifuge.
8. Hemocytometer.
9. PBS (Thermo Fisher Scientific, 10010-023).
10. Cell culture incubator capable of regulating temperature (37°C), humidity, and carbon dioxide (5%).
11. Water bath at 37°C.
12. Biosafety hood suitable for growing cells aseptically.

### 2.3 Expanding OP9 cells

1. OP9 cells (ATCC, CRL-2749)
2. OP9 growth medium: 500 mL Minimum essential medium a, no nucleosides (Thermo Fisher Scientific, 12561-056), 100 mL Fetal bovine serum (FBS; 20% final concentration, Thermo Fisher Scientific, 10437028), 5 mL penicillin/streptomycin (1% final concentration, Thermo Fisher Scientific, 15140-122).
3. 0.25% Trypsin-EDTA, phenol red (Thermo Fisher Scientific, 25200056).
4. T25 flasks, vented cap (Thermo Fisher Scientific, 12556009).
5. 12-well plates (Fisher Scientific, 12-556-005).
6. Cell culture incubator capable of regulating temperature (37°C), humidity, and carbon dioxide (5%).
7. Biosafety hood suitable for growing cells aseptically.

### 2.4 Differentiating mESCs

1. 12-well plate with OP9 cells ~80% confluent in each well.
2. 5,000-7,500 mESCs.
3. 0.25% Trypsin-EDTA, phenol red (Thermo Fisher Scientific, 25200056).
4. Recombinant murine thrombopoietin (TPO) (VWR, 10770-952).
5. PBS (Thermo Fisher Scientific, 10010-023).
6. Bovine Serum Albumin (Fisher Scientific, BP1600-100).
7. Cell culture incubator capable of regulating temperature (37°C), humidity, and carbon dioxide (5%).
8. Biosafety hood suitable for growing cells aseptically.

### 2.5 Docking transgenes into the genome of mESCs using φC31 integrase

1. Eight to twelve wells in a 12-well plate with confluent mESCs growing on mitomycin C treated MEFs (this can be scaled up or down, depending on the experimental design).
2. Lipofectamine 2000 (Thermo Fisher Scientific, 11668-019).
3. Opti-MEM (Thermo Fisher Scientific, 31985070).
4. Plasmid for inserting transgene into mouse genome: pBT378_pattB-pCA-GFP-pA-attB plasmid (Addgene, 52554).
5. Plasmid with φC31 integrase to enable recognition and integration at the *attP* location in the Hipp11 chromosome (Addgene, 13795).
6. Microcentrifuge tubes (Fisher Scientific, 05-408-129).
7. Biosafety hood suitable for growing cells aseptically.

### 2.6 Flow Cytometry analysis of differentiation and loaded transgene markers

1. PBS (Thermo Fisher Scientific, 10010-023).
2. Bovine serum albumin (BSA) (Fisher Scientific, BP1600-100).
2. Hoechst 33342 solution for identifying live cells (Fisher Scientific, BDB561908).
3. CD117 (c-Kit) clone 2B8 for identifying hematopoietic stem cell progenitors, PerCP conjugated (VWR, 105821).
4. CD45 for identifying nucleated hematopoietic cells, APC conjugated (30-F11) (VWR, 103111).
5. CD41 for identifying MKs and platelets, PE conjugated (Fisher Scientific, BDB561850).
6. Flow cytometry: Beckman Coulter Life Sciences CytoFLEX S cytometer.

### 2.7 Imaging for analysis of differentiation and loaded transgene markers

1. 12-well plates (Fisher Scientific, 12-556-005).
2. PBS (Thermo Fisher Scientific, 10010-023).
3. 4% formaldehyde (Thermo Fisher Scientific, FB002).
4. Permeabilizing blocking solution: 5% Normal donkey serum (Jackson ImmunoResearch Laboratories, 017-000-121), 1% Bovine Serum Albumin, IgG free, protease free (Jackson ImmunoResearch Laboratories, 001-000-162), 0.25% triton (Fisher Scientific, BP151-500).
5. Blocking solution: 5% Normal donkey serum (Jackson ImmunoResearch Laboratories, 017-000-121), 1% Bovine Serum Albumin, IgG free, protease free (Jackson ImmunoResearch Laboratories, 001-000-162).
6. Glycerol (Fisher Scientific, G31-1).
7. DAPI to stain nuclei (Fisher Scientific, PI62247).
8. Phalloidin to stain actin filaments (Thermo Fisher Scientific, A30106).
9. Rat anti-CD41 primary antibody, unconjugated to stain MKs and platelets (Thermo Fisher Scientific, MA516875).
10. Donkey anti-rat secondary antibody, Alexa Fluor® 647 AffiniPure Donkey Anti-rat IgG (H+L) (Jackson Labs, 712-605-150).
11. Mouse anti-GFP primary antibody, unconjugated to stain for docked GFP (Thermo Fisher Scientific, MA5-15256).
12. Donkey anti mouse secondary antibody, Alexa Fluor® 647 AffiniPure Donkey Anti-Mouse IgG (H+L) (Jackson Labs, 715-605-150).
13. Zeiss Axio Observer 7 live cell imaging inverted fluorescence microscope.
14. Nikon TS100 inverted microscope.

## 3. Methods

An overview of the entire differentiation process described in this protocol is shown in Fig. 1.

**Figure 1:**
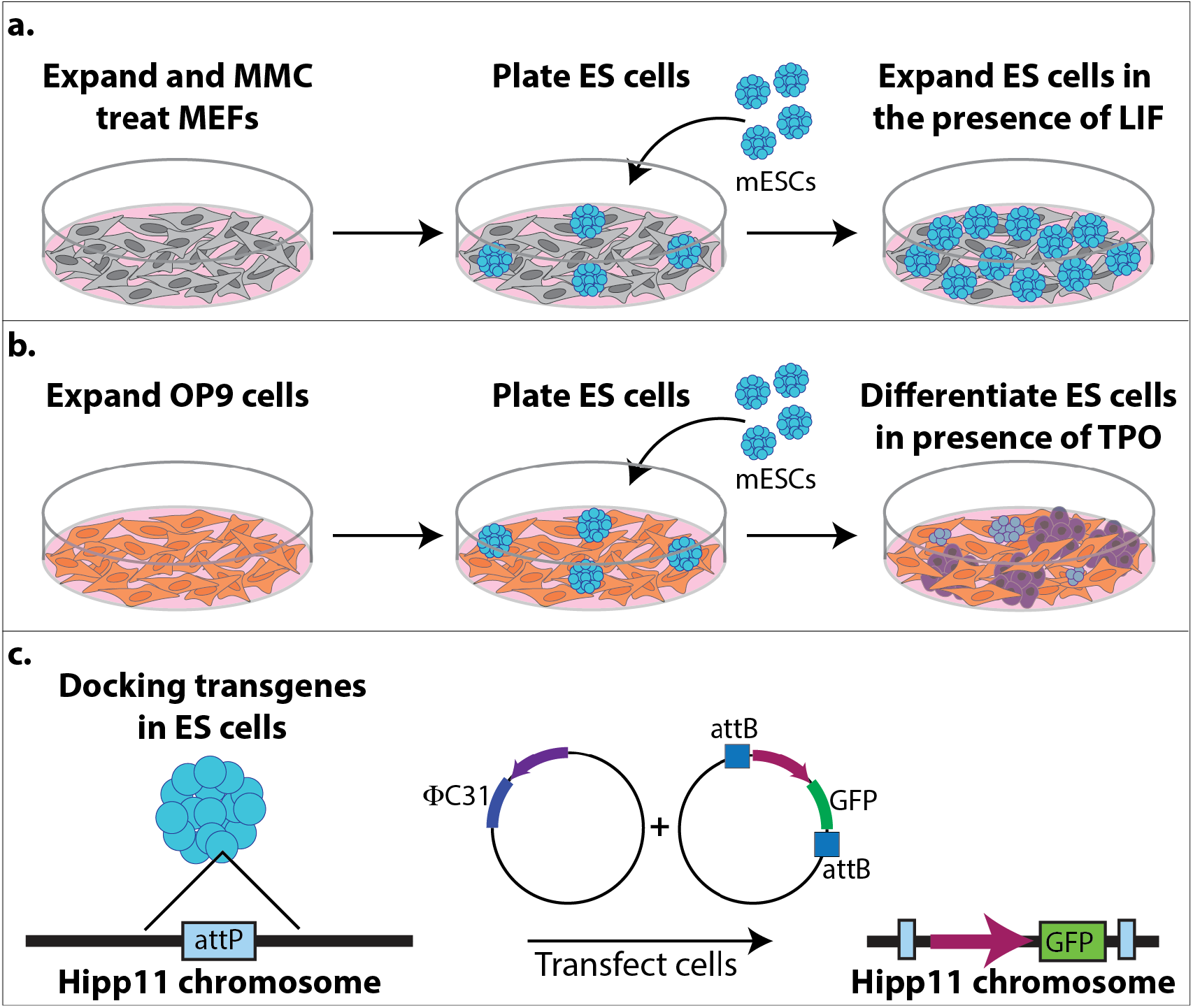
Schematic of the differentiation process. **(a)** Expansion and mitomycin C (MMC) treating mouse embryonic fibroblasts (MEFs, grey cells) for supporting the growth and expansion of mouse embryonic stem cells (mESCs, turquoise cells). **(b)** Expansion of OP9 cells (orange cells) for differentiating mESCs into megakaryocytes (MKs, purple cells) in the presence of thrombopoietin (TPO). **(c)** Docking transgenes in the Hipp11 chromosome of mESCs using φC31 integrase. The mESCs were harvested from a TARGATT mouse line that has an *attP* site located in the Hipp11 chromosome for φC31 integrase to catalyze the recombination and insertion of between *attB* and *attP* sites to insert a promoter and GFP into the Hipp11 chromosome.

### 3.1 Expanding and mitomycin C treating MEFs

1. Make a 0.1% gelatin solution (0.1g/100 mL Milli-Q water) by weighing out 0.1 g of gelatin and adding it to a beaker of 100 mL Milli-Q water with a magnetic stirrer and placed on a magnetic stirrer hotplate (*see* **Note 3**). Once the gelatin is completely dissolved, let cool to room temperature, then filter sterilize by running through a sterile disposable filter unit. Coat a T75 flask with 5 mL of 0.1% gelatin. Rotate flask to ensure the entire bottom of the flask is covered. Let sit at room temperature in the hood for 20-30 minutes.
2. Thaw a vial of the neomycin resistant MEFs by moving tube around in a 37°C water bath (*see* **Note 4**).
3. Once thawed, wipe water from tube and spray with 70% ethanol before placing in hood. Transfer cells to 7 mL of pre-warmed MEF growth media in a 15 mL conical tube (*see* **Note 5**).
4. Centrifuge cells at 300 xg for 5 minutes.
5. Aspirate medium off of cell pellet, being careful not to disturb the pellet.
6. Aspirate the gelatin off of the T75 flask (no need to wash the flask)
7. Resuspend the cells in 10 mL MEF growth medium and transfer to a T75 flask.
8. The next day change the medium to get rid of the cells that did not survive the freeze-thaw by aspirating medium off of cells and adding 10 mL of fresh pre-warmed MEF growth medium (*see* **Note 5**).
9. Check on cells daily to ensure they are growing (Fig. 2a and b). About 1-2 days after thawing, when cells are about 90% confluent (Fig. 2c), pass into 2x-T175 (or 5x-T75 flasks) gelatin coated flasks (*see* **Note 6**).
10. Once the cells are ready for passage (Fig. 2c), pass cells into 10x T175 gelatin coated flasks (*see* **Note 6**).
11. When cells are about 90% confluent (4-5 days), mitomycin C (MMC) treat the cells. MMC comes in 2mg of powder. Resuspend MMC in 2 mL sterile PBS to make it 1mg/mL (*see* **Note 7**). Add 100 μL of 1 mg/mL to 10 mL MEF growth medium so the final concentration is 10 μg/mL of MMC on the cells (*see* **Note 8**). Incubate the cells in the incubator for 2-4 hours (*see* **Note 9**).
12. Wash the cells with PBS by aspirating the medium with MMC off of the cells and adding 15 mL of PBS. Rotate the flask to ensure PBS is washed over all of the cells.
13. Aspirate the PBS and add 7 mL trypsin to each flask. Return to the incubator for about 5 minutes.
14. Check that the cells have detached in all flasks and once they have, add 14 mL MEF growth medium and transfer all cells to 50 mL conical tubes.
15. Count the number of cells using a hemocytometer.
16. Centrifuge cells at 300 xg for 5 minutes
17. Aspirate medium off of cell pellet, being careful not to disturb the pellet.
18. Resuspend cells in freeze medium (90% FBS, 10% DMSO) to freeze cells at 750,000-900,000 cells per cryovial (*see* **Note 10**).
19. Store at −80C overnight and transfer to liquid nitrogen the next day.

**Figure 2:**
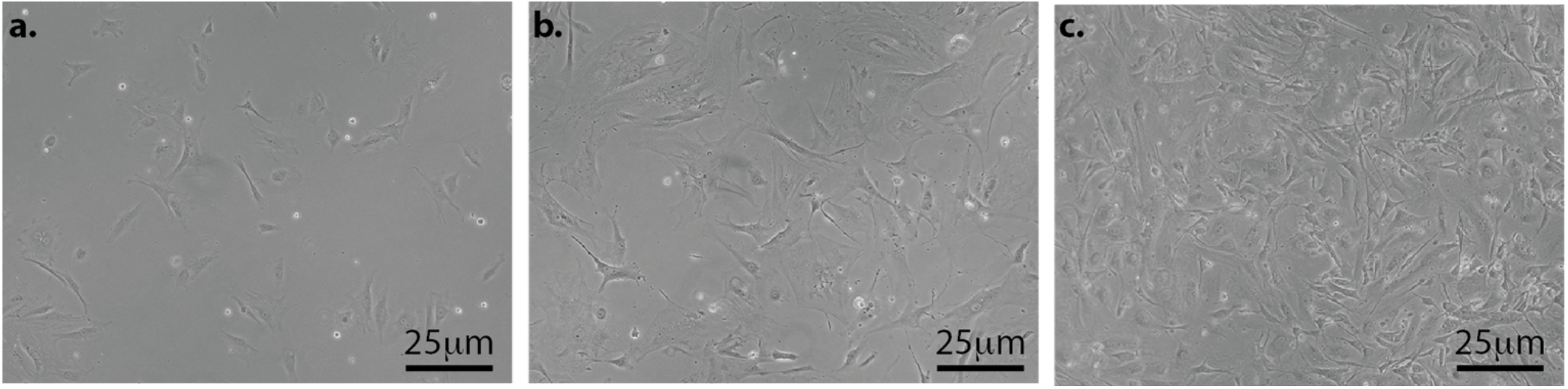
Bright field images of MEFs grown on gelatin coated plates. **(a)** and **(b)** are MEFs growing and not ready to be passed. **(c)** The cells have reached ~90% confluency and are ready to be passed.

### 3.2 Expanding mouse embryonic stem cells (mESCs)

1. The <u>day before</u> thawing mESCs, gelatin coat the bottom of a T25 flask with 2 mL of 0.1% gelatin. Rotate flask to ensure the entire bottom of the flask is covered. Let sit at room temperature in the hood for 20-30 minutes.
2. Thaw one vial of MMC treated MEFs by moving tube around in a 37°C water bath.
3. Once thawed, wipe water from tube and spray with 70% ethanol before placing in hood. Transfer cells to 9 mL of pre-warmed MEF growth media in a 15 mL conical tube (*see* **Note 5**).
4. Centrifuge cells at 300 xg for 5 minutes.
5. Aspirate medium off of cell pellet, being careful not to disturb the pellet.
6. Aspirate the gelatin off of the T25 flask (no need to wash the flask).
7. Resuspend the MEFs in 5 mL MEF growth medium and transfer to the T25 flask.
8. Place the flask in incubator overnight.
9. The next day, thaw a vial of the TARGATT mESC line by moving tube around in a 37°C water bath.
10. Resuspend the mESCs in 5 mL mESC growth medium with LIF and transfer to the T25 flask with the MMC treated MEFs that were thawed the previous day (*see* **Note 11**).
11. mESCs need fresh medium every day (*see* **Note 12**). To change the medium, aspirate the medium out of the T25 flask and replace with 5 mL pre-warmed ES medium containing LIF.
12. Carefully watch the growth of the mESCs daily (Fig. 3). The mESC colonies will grow over days and the day before they become very confluent (Fig. 3b), prepare a T25 flask of MMC treated MEFs (step 1) (*see* **Note 13**). The next day, the mESCs will be confluent (Fig. 3c) and need to be passed. Pass the cells 1:3 if you are actively running experiments or up to 1:25 if you need more time between passages (*see* **Note 6**).

**Figure 3:**
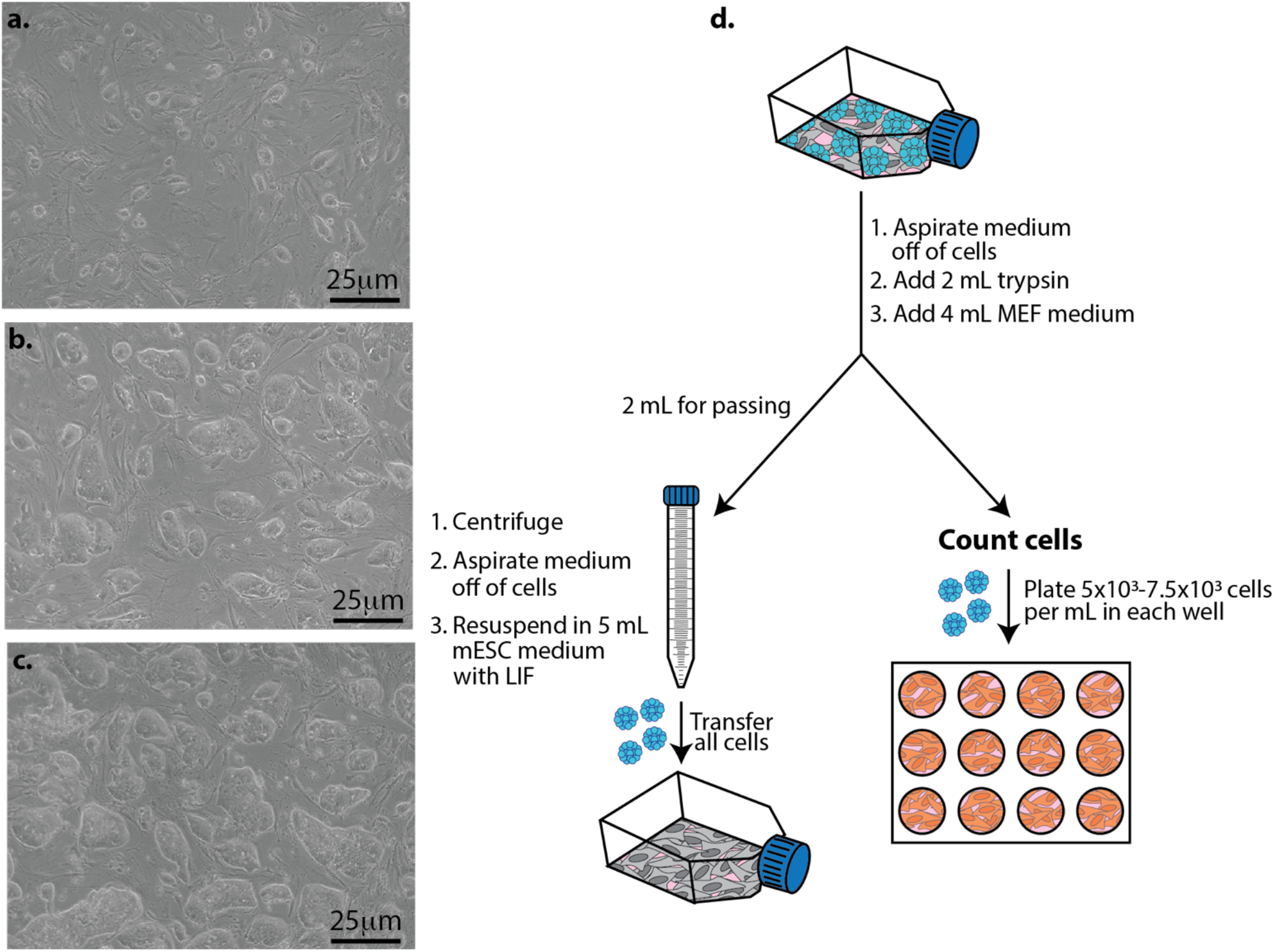
Growing mESCs on a layer of mitomycin C treated MEFs and plating mESCs for differentiation studies. **(a).** Representative bright field image two days after plating mESCs on MEFs. **(b)** Representative bright field image four days after plating mESCs on MEFs. **(c)** Representative bright field image five days after plating mESCs on MEFs. **(d)** mESCs (blue cells) are grown in a T25 flask until they are ready to pass (Fig. 3c). Pass cells into a T25 flask with MMC treated MEFs (grey cells) for expansion (left) and plate cells into a 12-well plate containing OP9 cells (orange cells) for differentiation studies (right).

### 3.3 Expanding OP9 cells and plating for mESC differentiation studies

1. Thaw a vial of OP9 cells by moving tube around in a 37°C water bath.
2. Once thawed, wipe water from tube and spray with 70% ethanol before placing in hood. Transfer cells to 7 mL of pre-warmed MEF growth medium in a 15 mL conical tube (*see* **Note 5**).
4. Centrifuge cells at 300 xg for 5 minutes.
5. Aspirate medium off of cell pellet, being careful not to disturb the pellet.
6. Resuspend the OP9 cell in 5 mL OP9 growth medium and transfer to the T25 flask.
7. Place the flask in incubator overnight.
8. Pass the OP9 cells when they reach about 80% confluency (Fig. 4b), at a ratio of 1:4 or 1:5. It is critical to watch the growth of these cells because the cell density is important. If the cells are too sparse (e.g. below 4×10^3^ cells/cm^2^) they will senesce and never reach confluency. It is also critical to not let the OP9 cells become over 80% confluent because the cells will start to differentiate into adipocytes and deposit lipid droplets in their cytoplasm (Fig. 4c). When OP9 cells start to deposit lipid droplets, they are no longer able to support the maintenance or differentiation of hematopoietic cells (*see* **note 14**).

**Figure 4:**
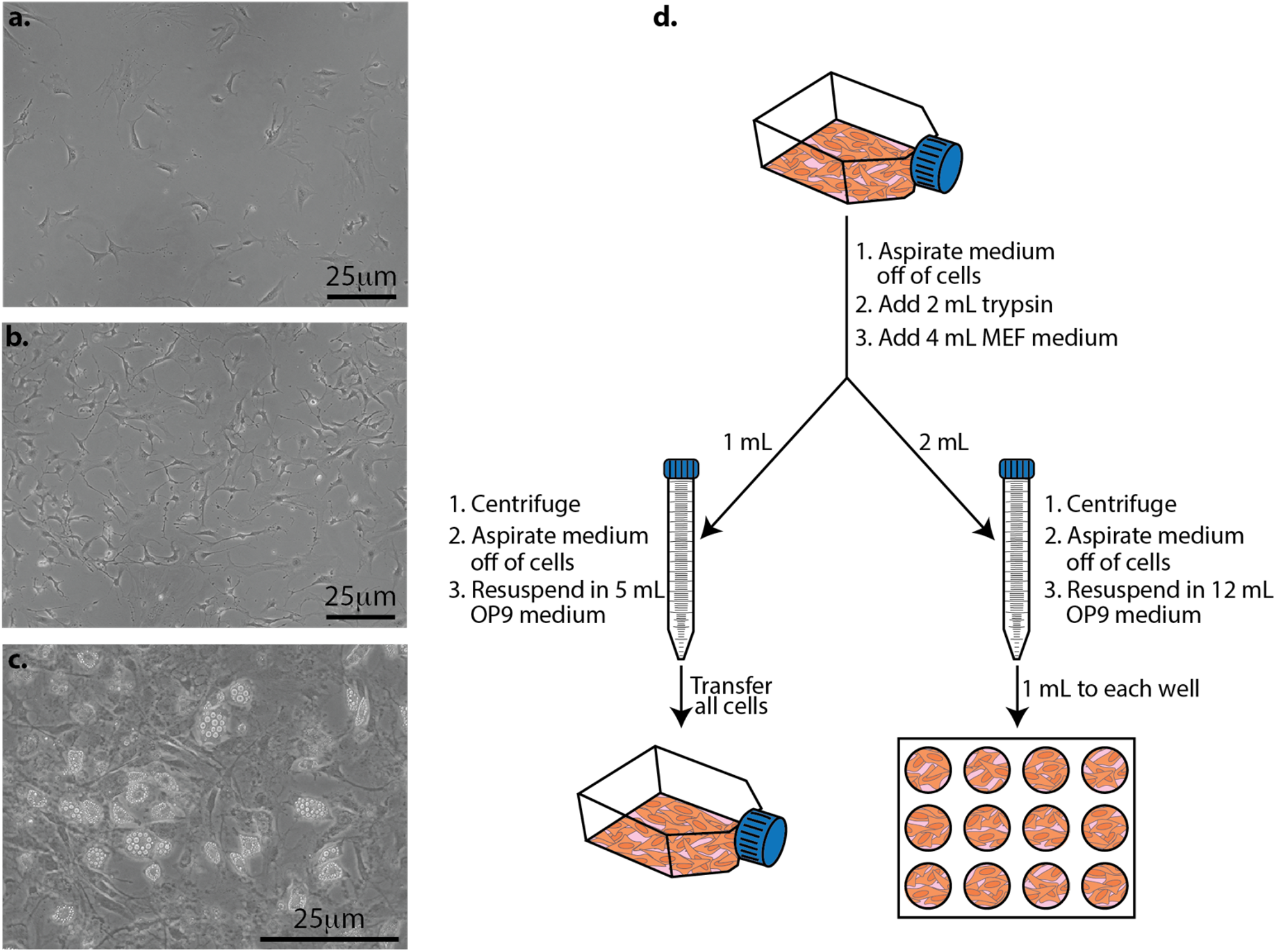
Expanding OP9 cells and preparing them for differentiation studies. **(a)** The day after passing OP9 cells. **(b)** Cells at ~80% confluency and are either ready for mESCs to be seeded on them for differentiation studies, or they need to be passed. **(c)** When cells become too confluent, they start to differentiate into adipocytes and deposit lipid droplets in their cytoplasm. At this stage, they will no longer support the maintenance and differentiation of hematopoietic cells and you should no longer use these cells. **(d)** OP9 cells (orange cells) are grown in a T25 flask until about 80% confluent (Fig. 4b). At this stage, the OP9 cells need to be passed into a T25 flask for expansion (left) and/or into a 12-well plate for differentiation studies (right).

### 3.4 Differentiating mESCs

#### 3.4.1 *Preparing OP9 cells* (Fig. 4)

1. You will need to prepare OP9 cells at least one day before the differentiation study can begin. When OP9 cells are ready to pass in a T25 flask (Fig. 4b), aspirate the medium out of the flask and add 2 mL of trypsin. Incubate at 37°C until the cells have detached from the flask, about 5 minutes.
2. Once the cells are detached, add 4 mL of complete MEF medium to deactivate the trypsin (*see* **Note 15**).
3. Add 1 mL of the resuspended cells to a 15 mL conical tube for passing (Fig. 4d, left), and 2 mL to a 15 mL conical tube for plating in a 12-well plate for differentiation studies (Fig. 4d, right). This can be scaled up or down depending on the scale of your differentiation studies. Discard the remaining cells.
4. Centrifuge cells at 300 xg for 5 minutes.
5. Aspirate medium off of cell pellet, being careful not to disturb the pellet.
6. Resuspend the pellet for passing in 5 mL OP9 growth medium and transfer to a T25 flask (Fig. 4d, left). Discard the remaining resuspended cells.
7. Resuspend the pellet for the differentiation study in 12 mL OP9 medium and plate 1 mL of the cells into each well of a 12 well plate (Figure 4d, right).
8. Place flask and plate in incubator.

#### 3.4.2 *Differentiating mESCs* (Fig. 5)

<u>Day 0</u>: Plate ES cells on prepared OP9 cells (Fig. 3d, right)
1. Aspirate medium off of a flask of confluent mESCs. Add 2 mL of trypsin and incubate at 37°C until the cells have detached from the flask, about 5 minutes.
2. Once the cells are detached, add 4 mL of complete MEF medium to deactivate the trypsin and transfer 1 mL of cells to a 15 mL conical tube (Fig. 3, left) (*see* **Note 16**).
3. With the remaining mESCs, count the number of cells using a hemocytometer.
4. Aspirate the OP9 medium off of the cells in the 12-well plate and add 1 mL of fresh OP9 medium to each well.
5. Plate between 5×10^3^ to 7.5×10^3^ mESCs per well in a 12 well plate. Because the volume of mESCs is so low (usually under 25 μL), you can add this small volume to each well without centrifuging. Indeed, some MEFs will be a part of this count, however, the number of mESCs is significantly higher than the number of MEFs during this passage so the MEFs are negligible. Going above the given range of mESCs will likely result in a failed differentiation study (*see* **Note 17**). Plating all 12 wells usually gives enough samples for assessing differentiation at different stages during the process, in addition to samples for imaging. You can scale up or down depending on the needs in your experiment.
6. Gently mix the mESCs in the fresh OP9 medium by rotating the plate front and back, then side to side, avoiding a circular pattern to prevent cells from pooling in the center of the wells.
7. Return the plate and T25 flask to the incubator until the next step.
<u>Day 3</u>: Half medium swap on differentiating cells.
1. Remove half of the medium from each well (500 μL) of the 12-well plate and add fresh OP9 growth medium (500 μL).
2. Return plates to the incubator until the next step.
<u>Day 4</u>: Pass OP9 cells (Fig. 4d)
1. Pass the OP9 cells according to your experimental set up and how many plates you will need for your differentiation study (see section 3.4.1).
<u>Day 5</u>: Pass differentiating mESCs (Fig. 5)
1. Transfer the medium growing on the cells to a 50 mL conical tube. This contains many semi- and non-adherent hematopoietic stem cells that you want to continue differentiating. It’s okay to pool all of the medium and cells from each well. Set aside (Fig. 5e, step 1).
2. To the remaining adherent cells in the 12-well plate, add 300 μL of trypsin to each well and return it to the incubator until the cells have detached (~5 minutes).
3. Once the cells have detached, add 600 μL MEF medium to each well to deactivate the trypsin. Gently pipette up and down each sample to separate any cell clumps. Return the plate to the incubator for ~20 minutes to allow the majority of the OP9 cells adhere to the plate (Fig. 5, step 2).
4. Collect the remaining suspension cells and pool them with the 50 mL tube of the semi- and nonadherent hematopoietic stem cells that you already collected in step 1 (Fig. 5, step 3).
5. Centrifuge cells at 300 xg for 5 minutes.
6. Aspirate medium off of cell pellet, being careful not to disturb the pellet.
7. Resuspend the pellet with 12 mL OP9 growth medium and add 20 ng/mL of TPO (*see* **Note 18**).
8. Take the plate of OP9 cells that was prepared the previous day and aspirate the medium off of the OP9 cells. Transfer 1 mL of differentiating mESCs in OP9 growth medium containing TPO onto the fresh plate of OP9s (Fig. 5, step 4).
9. Return plate to the incubator until the next step.
<u>Day 7</u>: Pass OP9 cells
1. Pass your OP9 cells according to your experimental set up and how many plates you will need for your differentiation study (see section 3.4.1) (Fig. 4d).
<u>Day 8</u>: Pass differentiating mESCs (Fig. 3d)
1. Repeat steps in Day 5 for passing differentiating mESCs (*see* **Note 19**).
<u>Day 12</u>: Assess suspension and adherent cells for differentiation (Fig. 6)
1. Assess differentiation using flow cytometry (section 3.6).
2. Assess differentiation using florescent microscopy (section 3.7).

**Figure 5:**
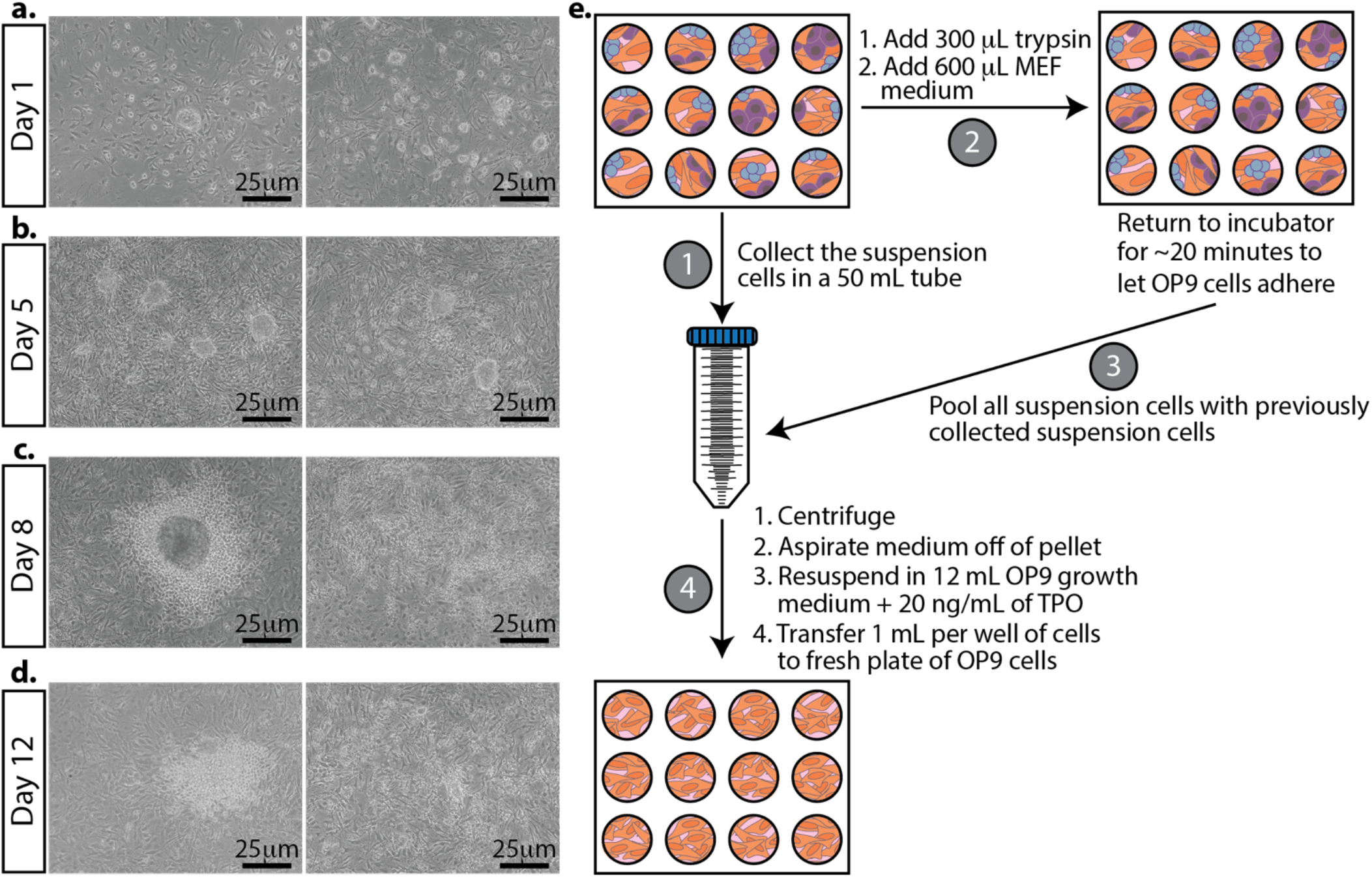
Morphological changes in mESCs during differentiation and passing differentiating mESCs. Representative bright field images. **(a)** The day after plating mESCs on OP9 cells. **(b)** mESCs grown on OP9 cells for 5 days. TPO is added at this stage. **(c)** Three days after adding TPO to the cultures. (**d)** The last day of differentiation. The cells are ready to be analyzed for differentiation markers. **(e)** Schematic for differentiating mESCs. First collect and pool the suspension cells from each well (left, labeled 1). Second, add trypsin to the adherent cells. Once the cells are detached and the trypsin is neutralized with medium, return the plate to the incubator to remove the majority of the OP9 cells (they will attach first) for about 20 minutes (top, labeled 2). After the majority of the OP9 cells have attached, take the cells in suspension and pool them with the previously collected suspension cells (right, labeled 3). Plate onto a fresh plate of OP9 cells growing in a 12-well plate (bottom, step 4).

**Figure 6:**
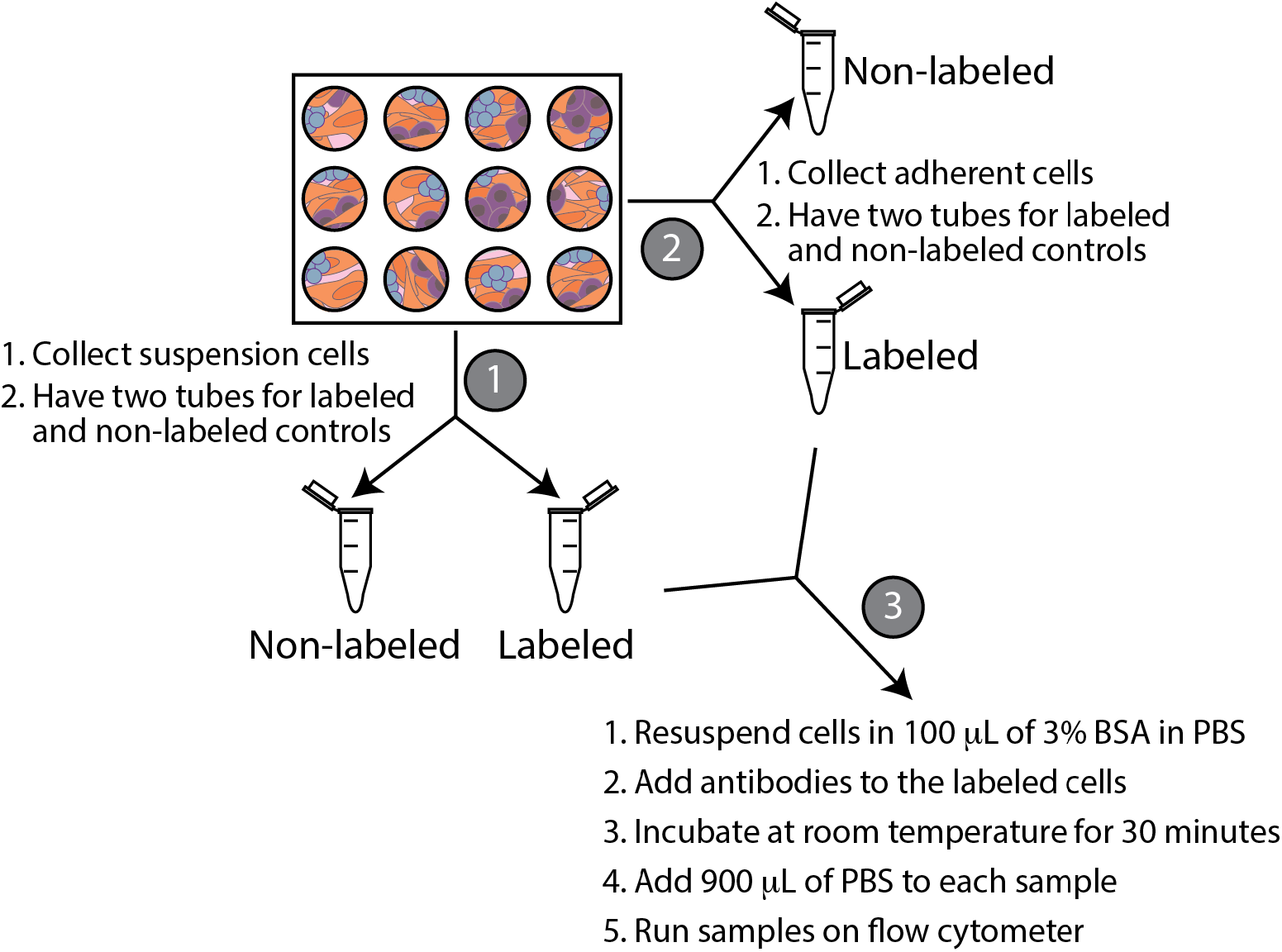
Schematic of collecting cells for flow cytometry. Collect the suspension cells and split them into two tubes (step 1). Next collect the adherent cells (step 2). Lastly, resuspend cells in 3% BSA and label the appropriate tubes with antibodies (step 3).

### 3.5 Docking transgenes into the genome of mESCs using fC31 integrase

1. Two days before starting the differentiation of transfected mESCs, prepare a 12 well plate of MMC treated MEFs (see section 3.2) (*see* **Note 20**).
2. The day before transfecting mESCs, when passing mESCs (Fig. 3d), count the number of mESCs and transfer 1×10^6^ to 3×10^6^ mESCs to a 15 mL conical tube.
3. Centrifuge cells at 300 xg for 5 minutes.
4. Aspirate medium off of cell pellet, being careful not to disturb the pellet.
5. Resuspend the pellet with 12 mL mESC growth medium with LIF.
6. Aspirate the MEF medium from the 12 well plate.
7. Distribute 1 mL of mESCs to each well of the 12 well plate with MMC treated MEFs.
8. Return plate to the incubator for a few hours to allow the mESCs to settle on the MEFs (*see* **Note 21**).
9. Later in the afternoon, after the mESCs have settled onto the MMC treated MEFs, gather all of the reagents for a Lipofectamine 2000 transfection. We are doing a co-transfection of φC31 (the integrase) and attB-GFP (the transgene to be inserted). Follow the manufacturer’s instructions for the Lipofectamine 2000 transfection. After transfecting the cells, grow them in mESC medium containing LIF overnight.
10. The next day, pass the transfected mESCs onto a prepared plate of OP9 cells to begin the differentiation process (section 3.4.2).

### 3.6 Flow Cytometry

1. We usually take 1 well of the 12 well plate for assessing differentiation at different time points using flow cytometry. Adjust as needed for your experiments.
2. Collect the suspension cells by pipetting the medium from one well of the 12 well plate into two microcentrifuge tubes (Fig. 6). One tube will be used as a non-labeled control and the other tube will be labeled with antibodies.
3. Collect the adherent cells by adding 300 μL of trypsin to the well that you already collected the suspension cells from. Once the cells are detached, add 600 μL of MEF growth medium to deactivate the trypsin. Add 450 μL of adherent cells to two microcentrifuge tubes. One tube will be used as a nonlabeled control and the other tube will be labeled with antibodies.
4. Add 5 μL of Hoechst per mL of growth medium to the microcentrifuge tubes that will be labeled with antibodies. Place the tubes in the incubator for 30-60 minutes, protected from light.
5. Centrifuge the labeled and non-labeled cells at 300 xg for 5 minutes.
6. Aspirate medium off of cell pellet, being careful not to disturb the pellet.
7. Resuspend cells in 100 μL of 3% Bovine Serum Albumin (BSA) in PBS (*see* **Note 22**).
8. Add 2 ng/μl of CD41 antibody, 2 ng/μl CD45 antibody, and 50 ng/μl of c-Kit antibody to the tube with the cells already labeled with Hoechst.
9. Incubate at room temperature for 30 minutes, protected from light.
10. Dilute samples with 900 μl of PBS.
11. Run samples on flow cytometer (Fig. 7).

**Figure 7:**
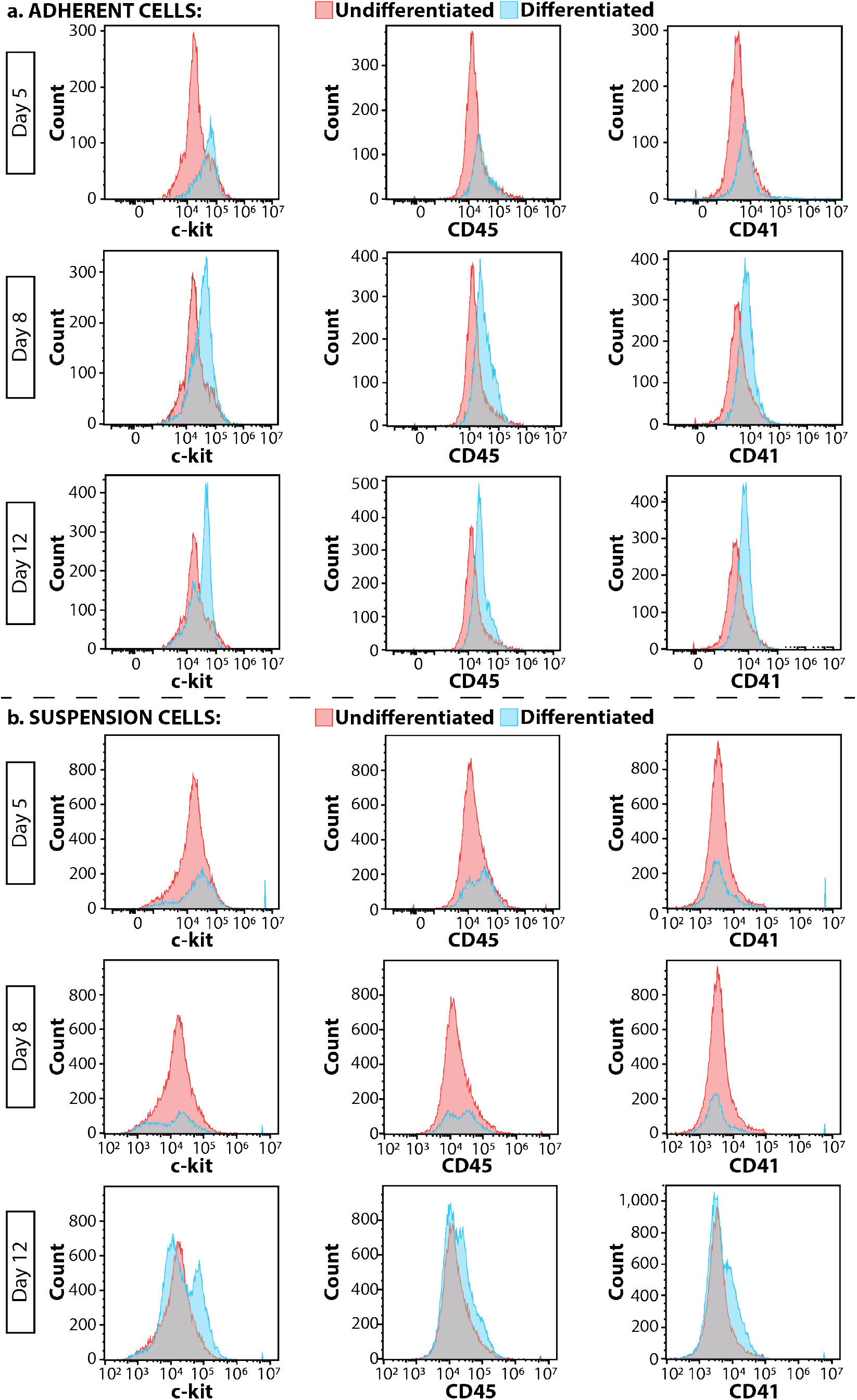
Flow cytometry to assess differentiation outcomes of the **(a)** adherent cells, and **(b)** the suspension cells. Pink peaks are undifferentiated TARGATT cells and blue peaks are differentiated cells.

### 3.7 Imaging

1. Aspirate medium off of cells you will be imaging and wash with cells with 1 mL PBS and aspirate it off of the cells (*see* **Note 23**).
2. Add 500 μL (enough to cover cells) of 4% formaldehyde. Incubate at room temperature for 10 minutes. Aspirate formaldehyde off of cells.
3. Wash cells with 1 mL PBS.
4. After aspirating the PBS off of the cells, add 500 μL of permeabilizing blocking solution. Incubate at room temperature for 10 minutes.
5. Wash cells with 1 mL PBS.
6. After aspirating the PBS off of the cells, add 500 μL of blocking solution. Add GFP primary antibody between at 1:500-1:2,000 dilution (we typically use 1:1000) and the CD41 primary antibody at 1:50-1:100 dilution (we typically use 1:100). Incubate at room temperature for 30 minutes.
7. Gently wash cells with 1 mL PBS two times.
8. After aspirating the PBS off of the cells, add 750 μL of blocking solution and add 0.5 μL of secondary antibodies and 0.75 μL of DAPI (*see* **Note 24**). Incubate at room temperature for 30 minutes.
9. Wash cells with 1 mL PBS and after aspirating the PBS add another 1 mL of PBS to keep the cells from drying out.
10. Image cells (Fig. 8).

**Figure 8:**
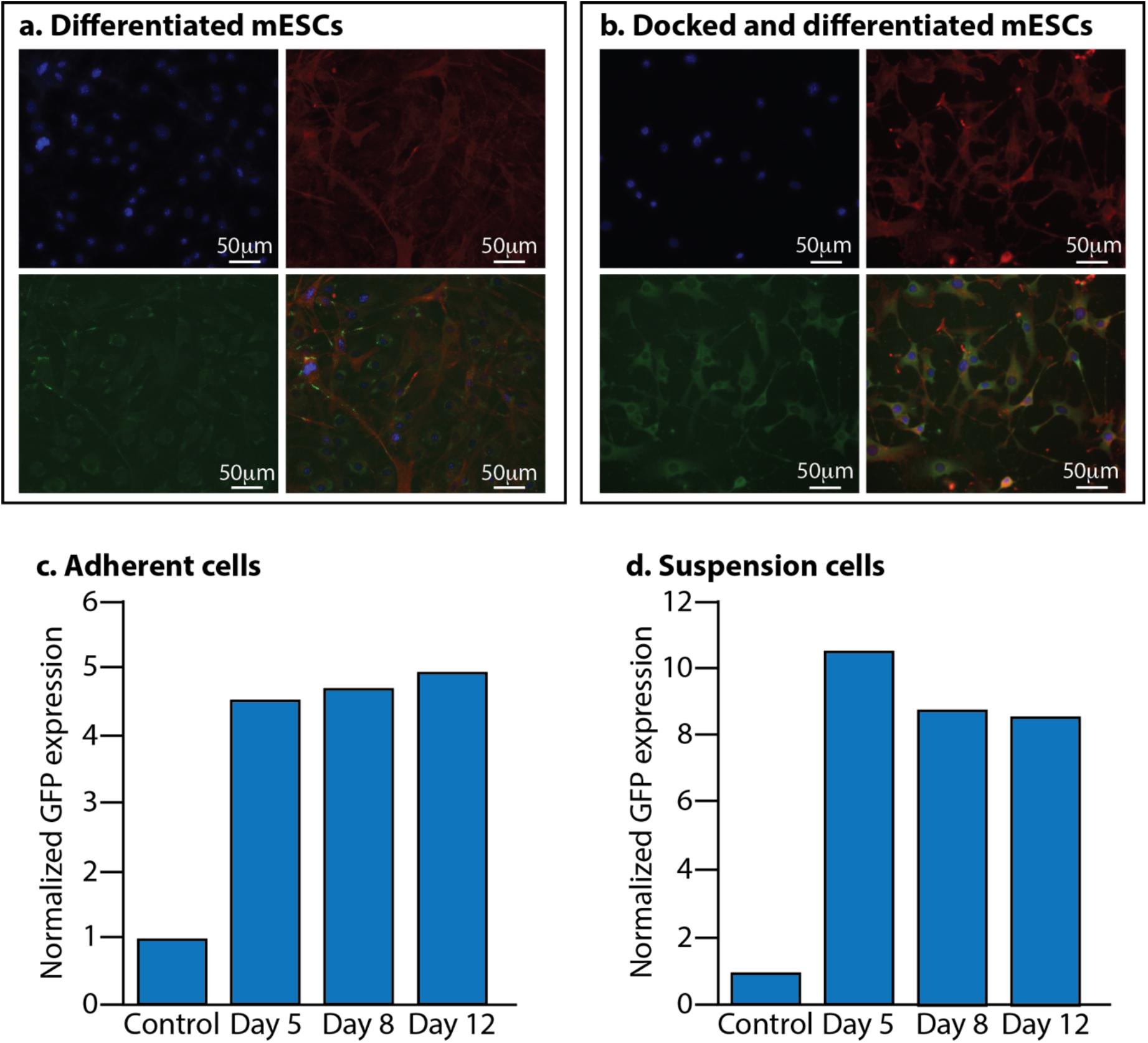
Assessing docked mESCs after differentiation. **(a)** Non-docked mESCs differentiated into MKs. Blue is DAPI, red is phalloidin, green is CD41. Bottom right is the merged image. **(b)** Docked mESCs with GFP and differentiated into MKs. Blue is DAPI, red is phalloidin, green is GFP and the bottom right image is the merged image. **(c)** GFP expression of adherent cells during the differentiation process. **(d)** GFP expression of suspension cells during the differentiation process. In both **(c)** and **(d)** the control is non-docked TARGATT cells.

## 4. Notes

1. We have used this protocol with multiple mESC lines with similar results.
2. LIF comes lyophilized at 10 μg. Resuspend in 100 μL sterile PBS. After making the complete ES medium without LIF, take a 50 mL aliquot and add 5 μL of the resuspended LIF. This can be stored at 4°C for up to a week. The LIF will degrade so only add it to smaller aliquots of mESC medium that will be used within a week.
3. Gelatin goes into solution faster with constant stirring and warming up the solution by using a magnetic stir bar with a magnetic stirrer hotplate. Larger batches of gelatin can be made and stored at 4°C for months. Always check that the gelatin is clear before using. If the solution looks cloudy it is likely contaminated. Throw away immediately and make fresh gelatin.
4. Neomycin resistant MEFs can be used when selecting for stable mESC lines containing a neomycin resistant cassette. We use these MEFs for the growth and expansion of all mESC lines that we grow, whether selecting for stable clones or not.
5. Pre-warm media in a 37°C water bath for about 10 minutes before using on cells. Once the media is warm, wipe off the water and spray entire bottle with 70% ethanol before placing it in the hood.
6. Passing cells: Add pre-warmed trypsin to cells (0.3 mL to each well of a 12-well dish; 2 mL to a T25 flask; 5mL to a T75 flask and 7 mL to a T175 flask). Always add double the amount of medium containing FBS to deactivate the trypsin once the cells have detached. Transfer cells to a 15 mL conical tube and centrifuge at 300 xg for 5 minutes. Aspirate the medium, taking care not to disturb the cell pellet. Resuspend the cells in fresh pre-warmed medium and transfer the cells to the appropriate growth vessel (5 mL for a T25 flask, 10 mL for a T75 mL flask, and 18 mL for a T175 flask).
7. Store any leftover MMC at −20C.
8. To conserve medium and FBS, transfer the medium from the growing cells (you will use about 13 mL per T175 flask) to 50 mL conical tubes (if you are growing 10x T175 flasks, you will transfer 130 mL of medium to 3x 50 mL conical tubes). Discard any remaining medium. Add 100 μL of 1 mg/mL of MMC to 10 mL media (e.g. 500 μL to 50 mL medium). Transfer 13 mL of medium + MMC to each flask. Rotate flasks to ensure all cells get covered with medium.
9. To fully MMC treat cells, incubate for no less than 2 hours and no more than 6 hours.
10. You can also freeze with 50% FBS, 40% MEF growth medium, and 10% DMSO, however, the higher FBS yields better survival after thawing.
11. ESCs grow better if they are plated on a layer of MMC treated MEFs that have settled in the flask. These cells are capable of supporting mESC growth for up to 10 days. In unusual circumstances when you need to pass your mESCs and you do not have a plate of MMC treated MEFs ready, you can thaw both the MEFs and the mESCs at the same time and plate them both in a gelatin coated T25 flask with mES growth medium containing LIF. If this method is used, it should be avoided for the next passage of the mESCs.
12. It is important that the mESC medium is changed daily to prevent mESCs from spontaneously differentiating. The only exception to this is that the day you pass the mESCs, you can double the medium (10 mL in a T25 flask) and not change the medium the next day.
13. Usually when you check on the mESCs to change the medium, the medium will start to turn orange in color when the cells are close to needing to be passed. Often the medium will turn yellow (and not cloudy) the next day when they need to be passed. It is important to take care of the cells (i.e. pass them) at this stage because the medium is more acidic and not ideal for the mESCs. If the cells are in this acidic medium too long, they will start to spontaneously differentiate.
14. As the OP9 cells undergo many passages, you will start to notice one or two cells with lipid droplets. These cells are usually okay to use in your studies, but you should consider thawing a fresh vial in the near future. Once you start to see the number of these cells with these lipid droplets, you should no longer use them for your differentiation studies.
15. Trypsin can be deactivated with double the volume of media with serum. For us, the base medium for making MEF growth medium is a lot less expensive compared to the OP9 medium. Therefore, our preference is to use MEF growth medium to deactivate the trypsin.
16. We use MEF growth medium to deactivate the trypsin with the mESCs because the FBS used in MEF medium is a lot less expensive than the FBS used to grow the mESCs.
17. The range of mESCs used in a differentiation study is critical for the differentiation study to progress to completion. We have found that adding more than 7.5×10^3^ of mESCs results in the OP9 cells becoming overwhelmed and they start peeling off of the plate, resulting in a failed differentiation attempt.
18. TPO comes 10 μg lyophilized. Resuspend in 1 mL PBS to give 10 μg/mL stock solution. For a concentration of 10 ng/mL in the differentiation medium, add 1 μL/mL of TPO to OP9 growth medium. For a concentration of 20 ng/mL in the differentiation medium, add 2 μL/mL of TPO to OP9 growth medium.
19. We have decreased the TPO concentration from 20 ng/mL to 10 ng/mL on day 8 with the TARGATT and other mESCs and our differentiation outcomes are similar to those reported here.
20. The surface area of a 12 well plate is about double to that of a T25 flask. If you plan on transfecting 12 wells, you will need 2 vials of the MMC treated MEFs since they are frozen at a density for a T25 flask. If you are only using 6 wells, you will only need 1 vial. MMC treated MEFs do not freeze-thaw well so once you thaw a vial of MMC treated MEFs, you should not freeze them again.
21. We usually pass the mESCs in the morning onto a plate of MMC treated MEFs, and in the afternoon do the transfection. You can let the mESCs grow overnight and do the transfection the next day. In both cases, the mESCs have a high rate of transfection.
22. To make 3% BSA in PBS, weigh out 3g of BSA and dissolve in 100 mL of PBS. Store at 4°C for short term and at −20°C for longer term storage.
23. We use a separate plate for imaging because the incubation steps are outside of the incubator and imaging the cells can take some time.
24. DAPI comes in 10mg, add 10 mL of MilliQ water to make 1 mg/mL working stock. This is 1000x so add 1μL to 1mL of blocking solution. The secondary antibodies come lyophilized. Resuspend these in a glycerol solution where you place a 50 mL conical tube on the scale and add 10 mL of MilliQ water. Tare the scale and drip 10 g of glycerol into the tube of water. Use this to resuspend the lyophilized secondary antibodies by adding 500 μL to each secondary antibody. Use this at 1:1500 to label cells.

## Acknowledgements

We would like to thank the generous support from the National Institute of Health Director New Innovator Award [1DP2CA250006-01]. Research reported in this publication was also supported by the Office of the Director of the National Institutes of Health under Award Number S10OD026959 and NCI Award Number 5P30CA042014-24.

